# The ultradian transcriptome of the human prefrontal cortex and abnormalities in schizophrenia

**DOI:** 10.1101/2022.05.26.493559

**Authors:** Madeline R. Scott, Wei Zong, Kyle D. Ketchesin, Marianne L. Seney, George C. Tseng, Bokai Zhu, Colleen A. McClung

**Affiliations:** Translational Neuroscience Program, Department of Psychiatry, Center for Neuroscience, University of Pittsburgh; Pittsburgh, Pennsylvania, USA; Department of Bioinformatics, University of Pittsburgh; Pittsburgh, Pennsylvania, USA; Aging Institute of UPMC, University of Pittsburgh School of Medicine; Pittsburgh, Pennsylvania, USA

## Abstract

Twelve-hour (12 h) ultradian rhythms are a well-known phenomenon in coastal marine organisms. While 12 h cycles are observed in human behavior and physiology, no study has measured 12 h rhythms in the human brain. Here we identify 12 h rhythms in transcripts which either peak at sleep/wake transitions (~9 AM/PM) or static times (~3 PM/AM) in the dorsolateral prefrontal cortex, a region involved in cognition. Subjects with schizophrenia lose 12 h rhythms in genes associated with the unfolded protein response and neuronal structural maintenance. Moreover, genes involved in mitochondrial function and protein translation, which normally peak at sleep/wake transitions, peak instead at static times in schizophrenia, suggesting suboptimal timing of these essential processes that may contribute to cognitive deficits commonly found in schizophrenia.

## Introduction

Twelve-hour (12 h) ultradian rhythms have long been observed in coastal marine animals, whose behavior align with ocean tides [1]. Recent studies have confirmed 12 h transcriptional rhythms in other organisms including *C. elegans*, mice, and olive baboons [1]. Various aspects of human behavior (sleep patterns, cognitive performance) and physiology (body temperature, blood pressure, migraine onset, circulating hormone levels) also exhibit 12 h rhythms [1]. However, as 12 h rhythms in transcript expression have not been identified in human tissue, it is unknown whether these processes are related to and/or regulated by molecular ultradian rhythms. Therefore, characterization of the human brain ultradian transcriptome will expand our understanding of transcript expression rhythms in the brain and their contribution to dysfunction in subjects with abnormalities in brain function.

Schizophrenia (SZ) is a chronic neuropsychiatric illness that affects over 20 million people worldwide and is a leading cause of disability [2]. Many SZ patients experience disturbances in the rhythmicity of sleep/wake cycles, peripheral gene expression, and daily hormones [3,4]. Molecular rhythm patterns, however, have only just begun to be directly measured in the human brain. To explore these rhythms in human postmortem brain tissue, our lab and others have utilized a “time of death” (TOD) analysis, in which gene expression data are organized across a 24 h clock based on the time of day of the subject’s death, to identify significant changes in gene expression rhythm patterns associated with specific brain regions [5], age [6], and psychiatric illnesses [7–9]. In SZ subjects, rhythmic analysis of RNA sequencing (RNA-seq) data collected by the CommonMind Consortium [10] from the dorsolateral prefrontal cortex (DLPFC) identified a loss of diurnal rhythmicity in a number of transcripts. Notably, SZ subjects exhibited 24 h rhythmicity in a set of transcripts which were not rhythmic in subjects with no psychiatric diagnosis (NP) [8]. Genes with enhanced 24 h rhythmicity in the SZ cohort were associated with mitochondria dysfunction and GABA-ergic signaling [8], consistent with previous work that finds differential expression of these pathways in subjects with SZ [11,12]. These studies demonstrate that circadian rhythms in gene expression can be reliably measured in human brain tissue and are severely disrupted in the DLPFC of SZ subjects. However, no study to date has attempted to measure 12 h rhythms in transcript expression in human brain or determine if there are changes to these ultradian rhythms in subjects with SZ.

In the current study, we use DLPFC data previously analyzed for circadian rhythms, allowing us to compare both 12 and 24 h rhythms within the same subjects. Multiple convergent analyses identify transcripts that have measurable 12 h rhythms in human DLPFC, with distinct abnormalities in the identity and timing of these transcripts in SZ.

## Results

### Multiple rhythmicity analyses identify 12 h rhythms in human DLPFC

We used a modified version of the nonlinear regression (NLR) TOD analysis used previously to determine circadian rhythms [5–9], in which a sinusoidal curve with a 12 h period is fit to gene expression across TOD, to measure 12 h rhythms in the human DLPFC of 104 subjects with known TOD (Fig 1A-D and S1A-B Fig; S1 Table). Out of the 13,915 detected transcripts, 819 (~6%) have significant 12 h rhythms at a threshold of p < 0.01, which we have previously used to delineate genes with 24 h rhythms in this cohort [8] (Fig 1E and S2 Table). As a confirmation of the NLR approach we also employed a Lomb-Scargle analysis, which finds the best fitting sinusoidal curve for each transcript in a manner that is unbiased in terms of period (S2A Fig) [13] and an eigenvalue/pencil analysis, which treats temporal gene expression as a composite rhythm and identifies four superimposed rhythmic components (RCs) that, when combined, best explain the temporal expression of each transcript (S2C-E and S3 Figs). In both the Lomb-Scargle and eigenvalue/pencil analyses we found enrichment of transcripts with 12 h periods (S2A, F Fig). 83% of transcripts identified as having 12 h rhythms in the NLR (p < 0.01) had a 12 h period in the Lomb-Scargle analysis (S2B Fig) and 68% had a 12 h RC in the eigenvalue/pencil analysis. Additionally, Ingenuity Pathway Analysis (IPA) found enrichment in mitochondria-related pathways (Oxidative Phosphorylation, Mitochondria Dysfunction, Sirtuin Signaling) for both NLR identified 12 h rhythms and eigenvalue/pencil identified 12 h RCs. The combination of these approaches gave us confidence that we had identified 12 h rhythms in the DLPFC.

**Fig 1.**
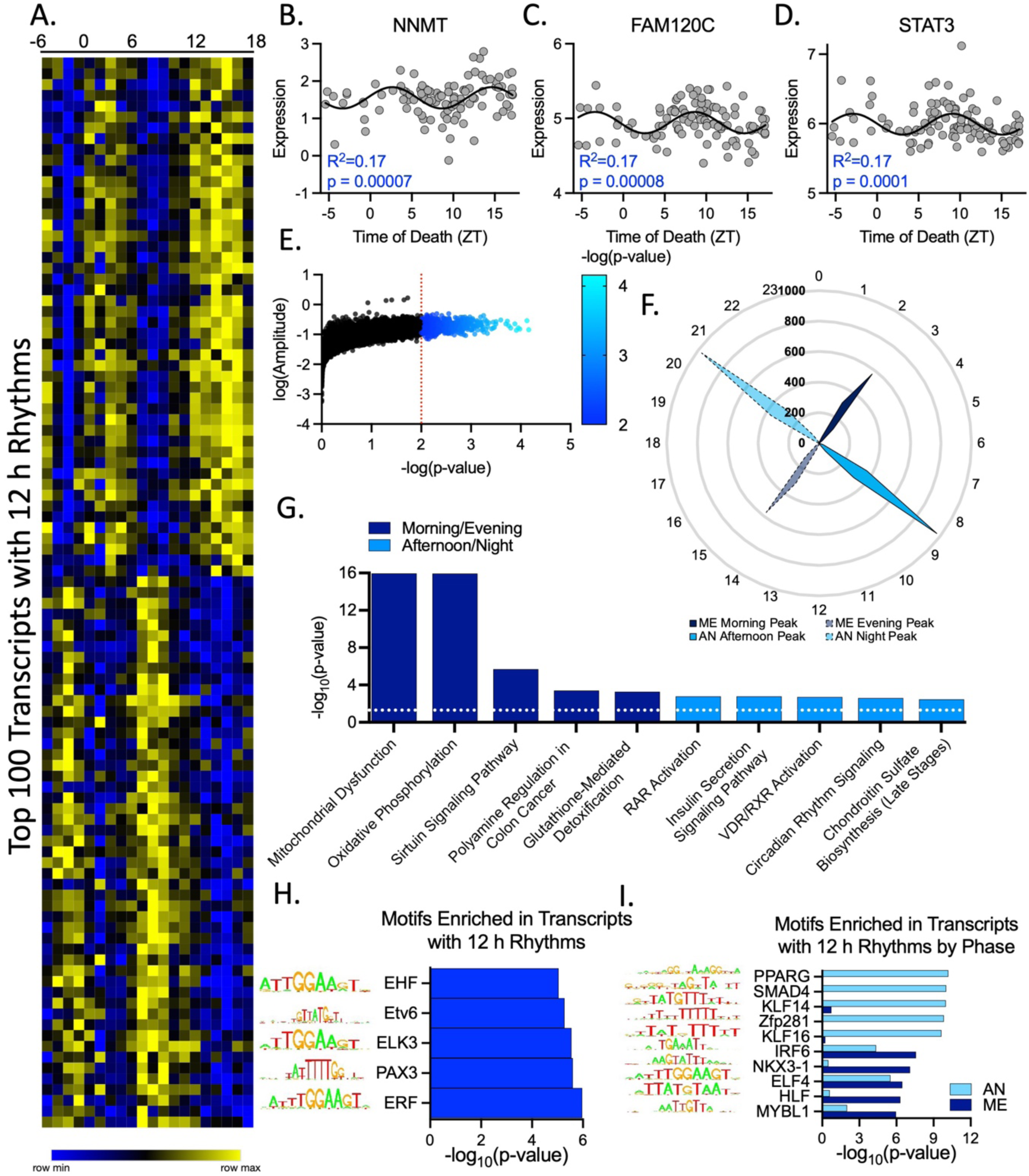
12 h rhythms in the human dorsolateral prefrontal cortex. (A-E) 12 h rhythms identified by a sinusoidal NLR. (A) Heatmap of top 100 transcripts with 12 h rhythms, ordered by phase. (B-D) Gene expression over TOD scatterplots for the top 3 transcripts with 12 h rhythms showing the sinusoidal curve, goodness-of-fit (R^2^), and p-value. (E) Total transcripts with significant 12 h rhythms in expression (p < 0.01, blue). (F) Radar plot showing peak expression times (ZT) of transcripts with 12 h rhythms. Solid lines indicate first peak, dashed lines indicated second peak. Morning/evening (ME) peak times indicated in dark blue, afternoon/night (AN) indicated in light blue. (G) IPA of transcripts with 12 h rhythms separated by peak time. (H-I) Motif enrichment analysis of transcripts with 12 h rhythms (H) all together and (I) separated by phase.

### Distinct timing patterns separate 12 h rhythms into two populations

The timing of 12 h rhythms revealed two distinct populations of transcripts, one which peaked in expression in the morning/evening (Zeitgeiber Time (ZT) 2-3 and 14-15; ~9 AM/PM) and the other that peaked during the afternoon/night (ZT 8-9 and 20-21; ~3 AM/PM) (Fig 1F). Transcripts associated with mitochondria and the proteasome (Polyamine Regulation in Colon Cancer) were enriched in the morning/evening population, while those associated with the cytoskeleton and calcium signaling (RAR Activation, Insulin Secretion Pathway, VDR/RXR Activation) peaked in the afternoon/night (Fig 1G).

We next determined potential sites of regulation and predicted upstream regulators for 12 h rhythms using a motif enrichment analysis [14] (Fig 1H-I). Motifs enriched in transcripts with 12 h rhythms are bound by transcription factors from the ETS domain family, Kruppel-Like Factors (KLF), and the SP domain family (Fig 1H and S4A-B Fig). Motifs associated with the ETS domain family remained enriched when we separately analyzed transcripts that peak in the morning/evening and afternoon/night, but both the KLF and SP domain families were associated only with motifs enriched in the afternoon/night (Fig 1I and S4C Fig). Motifs associated with the circadian-related Basic Helix-Loop-Helix (BHLH) domain family were not enriched in the analysis of transcripts with 12 h rhythms but were strongly enriched when analyzing the afternoon/night group separately (S4D Fig). Alternatively, motifs associated with PAR domain containing basic leucine zipper (bZIP) proteins, which have also been implicated in regulating circadian rhythms [15], were enriched both in the overall analysis (Fig 1H and S4A Fig) and in the morning/evening group (S4D Fig). Other families associated with motifs enriched in the morning/evening include the homeobox POU and PRD classes, while motifs associated with the SMAD family and peroxisome proliferator activated receptors (PPAR) were enriched in the afternoon/night group (Figs 1I and S4E Fig)

### Subjects with schizophrenia have fewer transcripts with 12 h rhythms

We next determined if 12 h rhythms were different in subjects with SZ, a psychiatric illness in which the DLPFC plays a central role [16]. Again, we used existing RNA-seq data produced from the CommonMind Consortium [10] (S1 Table and S1E-F Fig). We performed a sinusoidal NLR TOD analysis on a cohort of 46 SZ subjects and a group of 46 NP subjects taken from the full NP (fNP) cohort that best match the SZ cohort for TOD, sex, age, race, pH, and PMI (match NP (mNP); S1 Table and S1C-F Fig). Due to sample size limitations, we used a less stringent (p < 0.05) statistical cut off, consistent with our previous circadian analysis of these cohorts [8]. When we employed this cutoff, 1399 (10%) of transcripts had a 12 h rhythm in expression in the mNP cohort, while only 576 (5%) had a 12 h rhythm in SZ (Fig 2A-B, D). Of these transcripts, only 48 had a significant rhythm in both cohorts. A threshold-free approach, rank-rank hypergeometric overlap (RRHO) [17], supported our findings (Fig S5). Despite the lack in direct overlap, EIF2 signaling and mitochondria-associated pathways were the top pathways associated with transcripts showing 12 h rhythms for both the mNP and SZ cohorts (Fig 2C). Notably, only mNP subjects had 12 h rhythms in transcripts associated with the unfolded protein response (UPR) and RhoA Signaling (Fig 2C).

**Fig 2.**
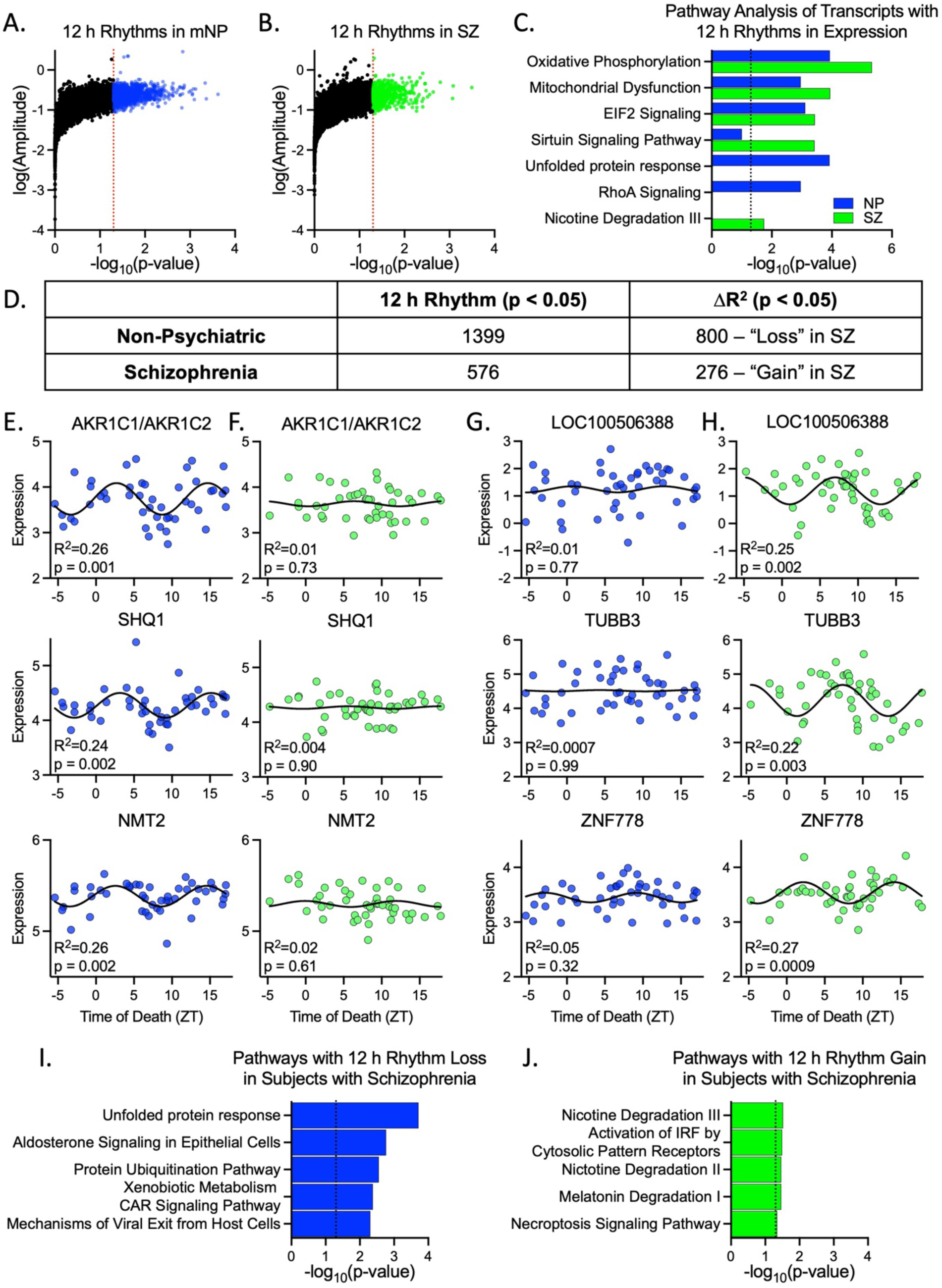
12 h rhythm reprogramming in schizophrenia. NLR analysis identified 12 h rhythms in the (A) mNP and (B) SZ cohorts. (C) IPA of transcripts with significant 12 h rhythms. (D) Number of transcripts identified as having a significant 12 h rhythm or a significant difference in goodness of fit (R^2^) between the two cohorts (deltaR^2^). (E-H) Examples of genes that (E-F) lose or (G-H) gain 12 h rhythms in SZ. Gene expression over time is shown for both the (E, G) mNP and (F, H) SZ cohorts. (I-J) IPA of transcripts that significantly (I) lose or (J) gain rhythmicity in SZ.

We next performed a loss/gain analysis of 12 h rhythmicity in SZ, as described previously for 24 h rhythms [8], to confirm our finding of fewer 12 h rhythms in SZ and to determine if transcripts experience ultradian reprogramming in SZ. 800 transcripts significantly lost 12 h rhythmicity and 276 transcripts gained rhythmicity in SZ (Fig 2D-H). Overall, transcripts that lost rhythmicity were associated with the UPR and the Protein Ubiquitination Pathway, while the smaller number of transcripts that gained rhythmicity did not fall into clear pathways (Fig 2I-J).

### 12 and 24 h rhythms converge on mitochondria-associated pathways in schizophrenia

Our previous work found a surprising gain of 24 h rhythmicity in mitochondria-related transcripts in subjects with SZ compared to the NP group [8]. Here we expanded upon those results using Metascape [18] to determine the biological processes implicated in the top transcripts with significant 12 and 24 h rhythms in mNP subjects and subjects with SZ (Fig 3A and S6 Fig). While each group had unique aspects of biological process enrichment, mitochondria and translation associated biological processes were enriched for both 12 and 24 h rhythms in SZ, but only 12 h rhythms in the mNP cohort (Fig 3A and S6 Fig).

**Fig 3.**
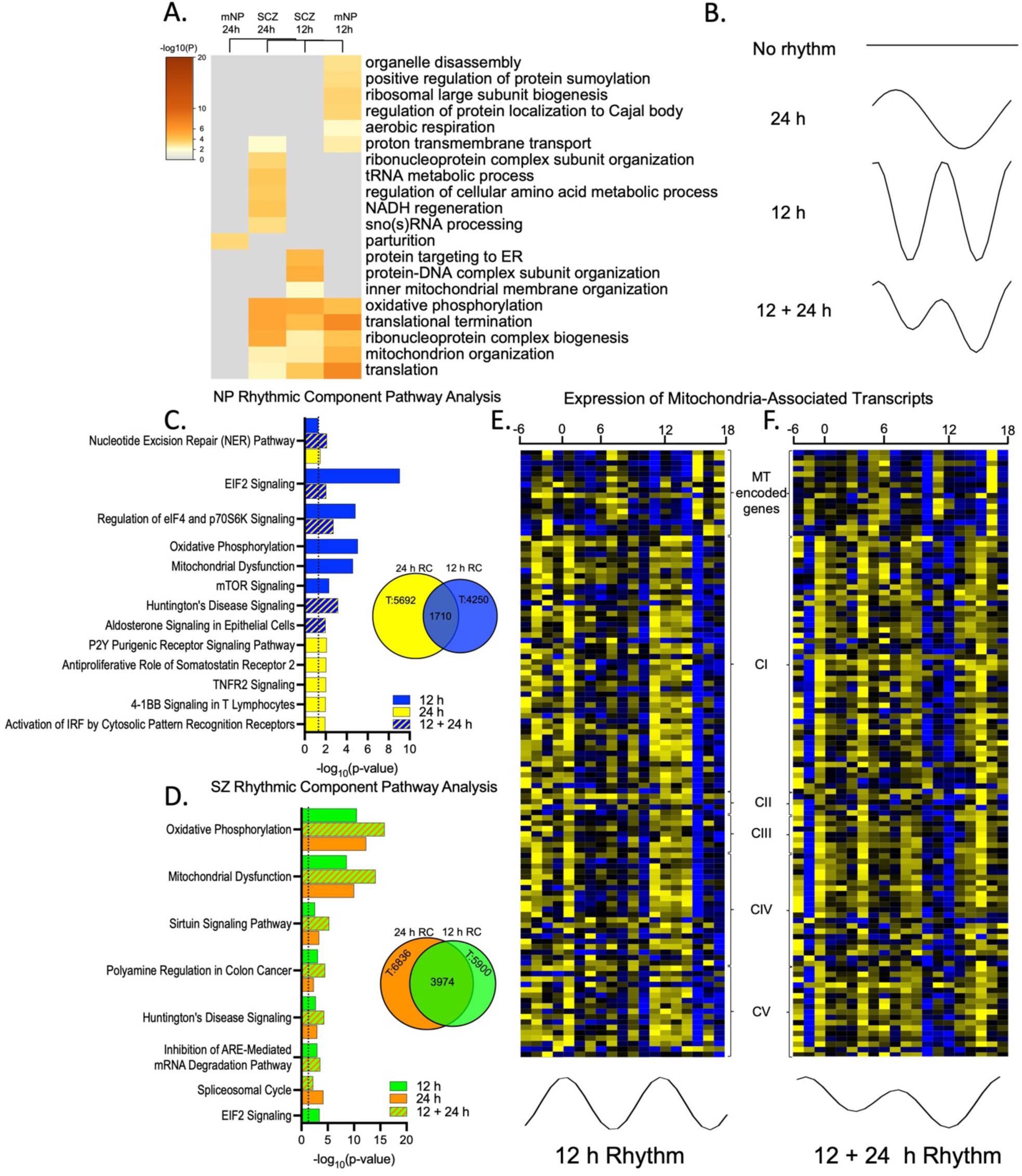
12 and 24 h rhythms converge on mitochondria associated transcripts. (A) Metascape-derived heatmaps of biological processes enriched for 12 and 24 h rhythms in mNP and SZ cohorts. (B) Illustrations of how transcripts could have no rhythms, a 24 h rhythm, a 12 h rhythm, or a combined 12 and 24 h rhythm. (C-D) IPA of transcripts with 12 h and 24 h RCs in (C) mNP subjects and (D) subjects with SZ. (E-F) Gene expression heatmaps of subunits from the mitochondria (MT) electron transport chain complexes (CI-CV) across time of day from (E) mNP subjects, which show a 12 h rhythm, and (F) subjects with SZ, which have a 12 + 24 h rhythm.

We next utilized the eigenvalue/pencil analysis – which allowed us to identify transcripts with gene expression that was best explained by either a single 12 h RC, a single 24 h RC, or a combination of both 12 and 24 h RCs (Fig 3B). Notably, the eigenvalue/pencil analysis identifies RCs – not overall rhythms in expression – and does not incorporate p-values [19]. This results in far more transcripts identified as having a 12 h RC by the eigenvalue/pencil analysis than a 12 h rhythm by the NLR analysis. Despite this, a very similar pattern in pathway enrichment emerged between the two analyses. In the mNP cohort, transcripts with 12 and 24 h RCs were found in distinct pathways (Fig 3C), while both 12 and 24 h RCs were linked to mitochondria-related pathways in SZ (Fig 3D). Intriguingly, the number of transcripts with both 12 and 24 h RCs increased from 1710 (30 and 40% of 24 and 12 h RCs, respectively) in the mNP cohort to 3974 (58 and 67% of 24 and 12 h RCs, respectively) in the SZ cohort (Fig 3C-D). As an example of what these data show, expression of transcripts in the electron transport chain complexes across time indicated mitochondria-associated genes had 12 h rhythms in the mNP cohort, while in SZ these transcripts had a combination of 12 and 24 h RCs (Fig 3E-F). We propose that this may explain why the NLR analysis identifies this group of transcripts as either a 12 h or 24 h rhythm, or both, in SZ subjects.

### Altered timing of transcripts with 12 h rhythms in schizophrenia

The SZ and mNP cohorts had similar timing patterns for transcripts with 12 h rhythms, with one population of transcripts that peaked in expression in the morning/evening, and the other that peaked during the afternoon/night (Fig 4A). However, IPA and Metascape analyses indicated that mitochondria, EIF2 signaling, and protein ubiquitination pathways were associated with morning/evening 12 h rhythmic transcripts in the mNP cohort, but with afternoon/night transcripts in the SZ cohort (Fig 4B,D and S7A Fig). Transcripts associated with the UPR also peaked during the morning/evening in the mNP cohort but, consistent with the loss of rhythmicity analysis, were not associated with either population in the SZ cohort (Fig 4B). Similarly, transcripts associated with the cytoskeleton and synaptogenesis (Synaptogenesis Signaling Pathway, Reelin Signaling in Neurons, Actin Cytoskeleton Signaling, RhoA Signaling) peaked in expression during the afternoon/night in the mNP cohort but were not associated with either timepoint in the SZ cohort (Fig 4B-C and S7A Fig). While most transcripts with 12 h rhythms peaked in the morning/evening in SZ, there was little distinct pathway recognition in either the IPA or Metascape analyses. In both analyses histone/chromosome regulation (histone h3 k36, positive regulation heterochromatin, regulation transcription initiation) was identified as a top pathway/biological process, though at a notably lower level of enrichment than the other groups (Figs 4B,E and S7B Fig).

**Fig 4.**
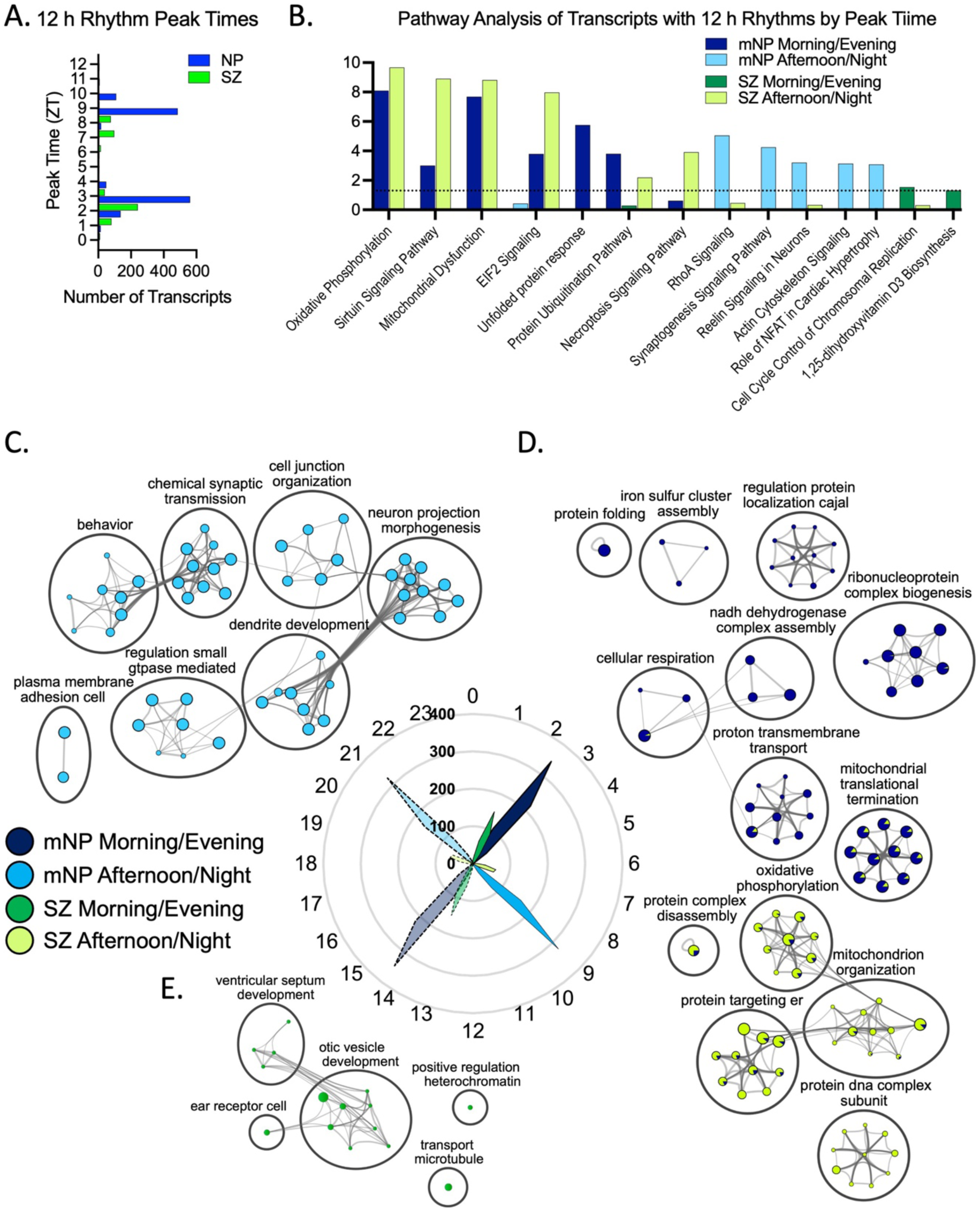
Altered timing of 12 h rhythms in schizophrenia. (A) Histogram of 12 h rhythm peak times in mNP and SZ. (B) IPA of transcripts with 12 h rhythms split by peak time. (C-E) Cytoscape visualization [20] of a Metascape analysis demonstrating the biological processes enriched for 12 h rhythms that peak during the morning/evening or afternoon/night.

### Multiple peak time-dependent patterns of motif enrichment differences in SZ

Finally, we applied motif enrichment analysis to these groups (mNP morning/evening, mNP afternoon/night, SZ morning/evening, SZ afternoon/night) to determine whether there may be differences in upstream regulators associated with 12 h rhythms in SZ (S8 Fig). Additionally, we split transcripts that significantly lost/gained 12 h rhythms by peak time and performed a motif enrichment analysis (S8 Fig). We found that ETS domain, PAR bZIP, POU class homeobox, PRD class homeobox, Forkhead box, and SRY-box families were associated with motifs enriched in the mNP morning/evening group of transcripts, but none of these families were enriched in the SZ morning/evening group (S8A-B, D Fig). Consistent with this, these families were associated with the morning/evening peaking transcripts that lost rhythmicity (S8A, C, G Fig). The ETS domain family, however, was enriched in the SZ afternoon/night group and in transcripts that gained rhythmicity in the afternoon/night, suggesting that, like the mitochondria pathways observed earlier, this protein family was associated with transcripts that have altered timing in SZ (S8A-B Fig). Similarly, the KLF, SP domain family, and BHLH domain family were associated with motifs enriched in transcripts that peak in the afternoon/night in the mNP cohort, but with the morning/evening group in the SZ cohort (S8A,F Fig). These distinct patterns in enrichment give us insight into which pathways may be regulating the multifaceted differences observed in 12 h rhythms in SZ.

## Discussion

Biological rhythms allow organisms ranging from bacteria to humans to anticipate changes in the environment across the light/dark cycle and adapt accordingly. These rhythms occur on many scales, from seasons to days to hours, and the importance of circadian rhythms in health and disease has become increasingly clear over the past few decades, particularly in the context of psychiatric illnesses [21,22]. Far less is known about ultradian rhythms, including how prevalent they are in the human brain, and whether they are disrupted in subjects with psychiatric disorders. In this study, we characterized the 12 h transcriptome within the DLPFC of NP and SZ subjects. These rhythms have distinct timing patterns, splitting them largely into two populations of transcripts. One of these peaks in the morning/evening (ZT 2-3/14-15; ~9 AM/PM) and is largely associated with mitochondria, EIF2 signaling, and the UPR. The other peaks during the afternoon/night (ZT 8-9/20-21; ~3 AM/PM) and is associated with cytoskeleton dynamics and the processes necessary to build and maintain neuronal connections.

These findings indicate that 12 h rhythms in the brain are associated with processes necessary for essential cellular functions – and may be fundamental for timing processes to maximize resources and reserve energy when not needed. This is consistent with previous analyses in mouse liver, which uncovered 12 h rhythms in metabolism-related transcripts and processes fundamental to transcription, RNA splicing, translation and proteostasis [23,24]. The timing of these processes coincided with sleep/wake transition times, leading to the proposal of a vehicle-cargo hypothesis for 12 h rhythms, in which 12 h rhythmicity accommodates increased demand for gene expression/processing at biological ‘rush hours’ (i.e. sleep/wake transitions) by upregulating expression of factors that facilitate protein and energy production [23]. Our observation that mitochondria- and translation-associated transcripts in human brain peak in expression in the morning/evening – when individuals are likely transitioning from wake to sleep or sleep to wake – strongly supports this hypothesis.

Strikingly similar to the mouse liver, our motif enrichment analysis implicated the ETS domain, KLF, and SP domain families in ultradian rhythm regulation (S4A Fig). Individual members of these transcription factor families also had significant 12 h rhythms, marking them as potential transcription factor regulators of these rhythms (S4B Fig). Consistent with our pathway analyses, separation of the data into two populations by peak time resulted in increased clarity. We found that transcripts which peak in the morning/evening are regulated by the POU class of homeoboxes and PAR bZIP family, while transcripts that peak in the afternoon/night are regulated by KLF, SP domain, and BHLH domain families (S4C Fig). The PAR bZIP and BHLH families are both implicated in circadian rhythms [15], suggesting that at least some proportion of what we have identified as 12 h rhythms could be regulated by the canonical circadian clock, however the BHLH family is very large and the circadian clock regulators are not the top hits within this group.

We observed far fewer transcripts with 12 h rhythms in SZ subjects than in the mNP subjects. Intriguingly, the UPR is the top pathway associated with transcripts that no longer have 12 h rhythms in SZ. The UPR has been shown to regulate 12 h rhythms in *in vivo* culture models and mouse liver [23,25]. Consistent with this, differential expression of UPR proteins and markers of UPR activity have been found in the DLPFC of subjects with SZ [26]. SZ subjects also do not display rhythmicity in transcripts associated with the actin cytoskeleton and synaptogenesis, which peak during the afternoon/evening in the mNP subjects (Fig. 4). Genomic analyses strongly implicate synaptogenesis and synaptic plasticity processes, while neuroanatomical studies have demonstrated reduced dendritic spine density in various regions, including the DLPFC, in SZ [27,28]. Of particular note, several voltage gated calcium channels (CACNAs), a couple of which are top SZ risk factor genes [27], are among this group with 12 h rhythms in mNP subjects but no rhythms in SZ.

Intriguingly, 12 h rhythms in mitochondria-associated pathways are not lost but instead switch from peaking during the morning/evening to afternoon/night timepoints in SZ. This could have profound effects on mitochondrial energy production at times of day when this is most needed. Various studies have implicated circadian rhythms in mitochondria biology, including gene expression in the mouse SCN, oxygen consumption and mitochondria respiration in isolated mitochondria from rat brains, and fusion/fission states in mouse macrophages [29–31]. Interestingly, mitochondrial-related pathways also gained 24 h rhythms in the DLPFC of subjects with SZ [8](Fig. 3), resulting in a convergence of rhythmicity dysregulation in mitochondrial function transcript expression. Gaining a 24 h rhythm may be a compensatory measure to account for sub-optimal timing of 12 h rhythms, or reflect changes associated with diminished neuronal activity specifically at night. Our findings are consistent with a robust literature of mitochondrial abnormalities in SZ, from genetics to function, number, location and shape [32]. It will be interesting in future studies to determine changes in mitochondrial morphology, number, function, and location in subjects with SZ as a function of time of day, and how this relates to transcript expression.

The superchiasmatic nucleus (SCN) is the master pacemaker of circadian rhythms in the brain, but whether ultradian rhythms are regulated across regions by a dedicated system is unknown. While not specific to 12 h rhythms, levels of dopamine in the striatum fluctuate in synchrony with ultradian locomotor activity cycles and dopaminergic transmission directly regulates ultradian cycle length in mice [33]. Interestingly, these dopamine-driven ultradian cycles in locomotor activity harmonize with circadian rhythms coordinated by the SCN, but if this relationship is disrupted it can lead to altered patterns of arousal and disrupted sleep/wake cycles when desynchronized [33], which are commonly observed in subjects with SZ. SZ has long been associated with altered dopaminergic transmission in cortico-striatal pathways [34]. Abnormalities in dopaminergic signaling, therefore, may be a potential mechanism by which ultradian rhythms are disrupted across the brain in SZ.

Human postmortem brain tissue research presents limitations that we have addressed as much as possible through our study design including exclusion of subjects older than 65, strict TOD criteria, and cohorts matched for a number of important biological, clinical and technical factors. Future studies in animal models will be necessary to determine the impact of features such as antipsychotic medication on the ultradian and circadian transcriptomes.

In conclusion, this study is the first to identify 12 h rhythms in transcript expression in the human brain. These rhythms are associated with fundamental cellular processes. However, in SZ, there is a strong reduction in the number of transcripts with 12 h rhythms, along with altered timing of transcripts important in mitochondrial function. Future studies will determine the functional consequences of these findings to optimal brain health and the pathophysiology of brain disorders.

## Materials and Methods

### Human postmortem brain samples

Human postmortem RNA-sequencing data for 613 samples were obtained from the CommonMind Consortium (https://nimhgenetics.org/available_data/commonmind/) and filtered based off previously described criteria [8,10]. Briefly, subjects were included if they met the criteria of rapid death (<2 h elapsed time between precipitating event and death announcement) and had a postmortem interval (PMI) of <30 h. In addition, subjects with age >65 years were removed as our lab has previously observed significant differences in molecular rhythms between young and elderly subjects [6]. 104 non-psychiatric (NP) subjects and 46 subjects with schizophrenia (SZ) met these criteria (S1 Table). The larger sample size of NP subjects may result in more statistical power for rhythm detection. Therefore, 46 non-psychiatric subjects that were best matched to the 46 SZ subjects by age, sex, race, TOD, PMI, site of collection, and pH (match NP cohort (mNP); S1 Table). The subjects in these cohorts are the same as those used in a previous study performed by our lab focused on circadian rhythms [8].

### Time of death analysis in the zeitgeber time scale

Prior to rhythmicity analysis, time of death (TOD) for each subject was normalized to a zeitgeiber time (ZT) scale as described in Seney et al.[8]. Briefly, the time of death for each subject was collected at local time then converted to coordinated universal time by adjusting time zone and daylight savings time. Coordinated universal time was further adjusted to account for longitude, latitude, and elevation of death place and each subjects TOD was set as ZT = *t* h after previous (if *t* < 18) or before next (if *t* ≥ −6) sunrise. The distribution of subject TODs is shown in S1 Fig.

### RNA-sequencing data preprocessing

Samples were analyzed using RNA-sequencing and 30,714 unique genes were identified. Genes were retained for analysis if counts per million (cpm) was greater than 1 in 50% of more subjects. All Y chromosome genes were eliminated along with transcripts with no identifiers. After filtering 13,914 genes remained, and expression of these genes were log2 normalized. Since samples in the CommonMind dataset were generated in 2 brain banks, we included equal proportions of Pittsburgh and Mt. Sinai individuals in each experimental group.

### Rhythmicity analyses

#### Nonlinear regression analysis

We utilized a modified version of the sinusoidal nonlinear regression analysis that has been used multiple times to detect circadian patterns of gene expression in human postmortem brain tissue (by our group and others) [5–9]. While previous papers defined the period of the sinusoidal curve as 24 hours, we modified this analysis to detect genes that best fit a sinusoidal curve with a 12 h period. P-values were obtained by comparing the observed coefficient of determination (R^2^) and a null distribution R^2^ that was generated by fitting the sinusoidal curve to 1,000 TOD-randomized data points. The results of this analysis can be found in S1 File.

#### Lomb-Scargle analysis

We performed a second analysis, the Lomb-Scargle periodogram, through the R package MetaCycle [13,35]. Lomb-Scargle is a method of rhythm detection primarily used in physics. It is unique in its capacity to analyze unevenly sampled time series data, identify rhythmic expression profiles with a range of periods, and to differentiate between periodic and non-periodic profiles for co-sine curves with high noise [13]. In this study, we analyzed the data for rhythms with periods between 8 and 28 h. These periods were chosen after testing several ranges and finding that there is an artificial build-up of rhythms with their period identified as the period boundary. 8 and 28 h appeared sufficiently far away from 12 and 24 h such that the shape of these transcript populations were unaffected by this artificial build-up. It is important to note when interpreting these results, that to get a true representation of transcripts with 24 h rhythms, we would have needed to use data with a 48 h TOD range. The results of this analysis can be found in S2 File.

#### Eigenvaluepencil analysis

Prior to analysis, weighted averages of every transcript were calculated for each hour interval for a range of ZT −5 – 42 (Fig. S2). Samples were then analyzed using the eigenvalue/pencil method as previously described [19,25]. This method assumes gene expression is the result of multiple superimposed oscillations and identifies the combination of rhythmic components that best explains the data without any constraints on the period, amplitude, or phase of the rhythms. The eigenvalue/pencil method was set to identify the top 4 rhythmic components (RC) contributing to gene expression. We used the eigenvalue/pencil method to identify four superimposed rhythmic components that best explain transcript expression for all 13,915 transcripts identified through RNA-sequencing in the DLPFC of non-psychiatric and schizophrenia subjects (Fig. S1). The period of these rhythmic components varied between 2 and Infinity (Fig. S2). The sampling rate of our cohorts, and the way in which we preprocessed data before applying the eigenvalue/pencil method, led us to consider periods below 9 or above 30 as indistinguishable from noise. The results of this analysis can be found in S3 File.

### Pathway enrichment and upstream regulator analyses

#### Ingenuity Pathway Analysis

Pathway and upstream regulator analyses were performed on transcript lists based on either the period (eigenvalue/pencil) or the p-value of the respective rhythmicity analysis. A user-provided list of the 13,915 transcripts expressed in the DLPFC was used as the background. 13,454 background transcripts remained, as some of the transcripts were not mapped or functionally annotated. Pathways and upstream regulators were considered enriched if they met a threshold of p < 0.05 (-log_10_(p-value) > 1.3 in figures). IPA pathways with < 15 or > 300 genes were not included in the analyses, and only direct relationships were considered for the upstream regulator analysis. The top 5 significant pathways and upstream regulators are represented for each analysis. Complete lists of all significant pathways and upstream regulators for each analysis can be found in S4 File.

#### Biological Process Enrichment

The web-based portal Metascape was used to perform biological process enrichment (May 2021) [18]. Process enrichment was accomplished using GO biological processes as the ontology source. Within Metascape, the multigene list meta-analysis option was used to compare process enrichment between 12 and 24 h rhythmic components or expression rhythms, diagnosis, and/or analysis method. Similar to IPA, transcript lists were based on either the period or the p-value (p < 0.05) of the respective rhythmicity analysis. However, Metascape analysis limits the number of transcripts to ≤ 3000. Only the eigenvalue/pencil analysis lists were longer than 3000 transcripts, so we used the top 3000 transcripts based on the largest amplitude values of the rhythmic components. A user-supplied list of the 13,915 transcripts expressed in the DLPFC was used as the enrichment background. 13,521 background transcripts remained, as some of the transcripts were not mapped or functionally annotated. Terms with p < 0.01, a minimum count of 3, and an enrichment factor of >1.5 were grouped into clusters based on their membership similarities (kappa score > 0.3). P-values were calculated based on the cumulative hypergeometric distribution and q-values were calculated based on the Benjamini-Hochberg procedure [18]. The most statistically significant term within a cluster was chosen to represent the cluster. The top 20 significant clusters of processes are depicted by Cytoscape [20]. If more than 10 enriched terms within a cluster were identified, the top 10 significant terms were chosen for visualization in the Cytoscape networks. Complete lists of all the enriched processes within each cluster can be found in S5 File. In the network plots, the nodes are represented as pie charts, where the size of the pie is proportional to the total number of gene hits for that specific term. The pie charts are color-coded based on the identity of the gene list, where the size of the slice represents the percentage of genes for the term that originated from the corresponding list. Terms that are similar (kappa score > 0.3) are connected by edges.

### LISA Cistrome Combined Motif Enrichment Analysis

The web-based LISA program was used to perform motif enrichment analyses [14]. For supplied gene lists, LISA determined whether a set of known motifs were enriched. The combined enrichment scores (-log10(pvalue)) were used to interpret results and can be found in S6 File.

### Data Visualization

#### Scatter Plots

We generated scatter plots of individual transcript expression for the top three transcripts with 12 h rhythms in expression. Each dot represents a subject with the *x*-axis indicating the TOD on ZT scale and *y*-axis indicating gene expression level after log2 normalization. The line superimposed on the graph is the fitted sinusoidal curve (Fig. 3A-C).

#### Heatmaps

Heatmaps for transcripts transcript expression across TOD were generated for the top 100 genes with significant 12 h rhythms in the fNP cohort (p < 0.01; Fig 1A) and for mitochondria electron transport chain complex genes in both the mNP and SZ cohorts (Fig 3E-F) using the matrix visualization and analysis software Morpheus (https://software.broadinstitute.org/morpheus). Expression levels were *Z-*transformed for each transcript and transcripts were ordered by the time at which they peak. Subjects were ordered by their TOD and grouped by ZT hour. Each column is the median gene expression for subjects within each ZT hour group.

#### Rank Rank Hypergeometric Overlap (RRHO)

RRHO is a threshold-free approach that identifies the overlap between two lists of transcripts ranked by their log_10_(p-value) [17]. This approach avoids an arbitrary threshold in conventional Venn diagram approaches.

## Acknowledgments

We would like to thank the Common Mind Consortium for providing the RNA sequencing data. In particular we thank Harry Haroutunian, Pamela Sklar, and Shaun Purcell for providing the time and date of death information for the Mt. Sinai samples. We also thank our brain bank at the University of Pittsburgh, primarily Jill Glausier and Carol Sue Johnston for time and date of death information for the Pitt samples. We also thank David Lewis and our members of the translational neuroscience program for helpful discussions. Most importantly we thank the family members that donated tissue.

## Author contributions

Conceptualization: MRS, BZ, CAM

Methodology: BZ, GCT

Investigation: MRS, BZ

Visualization: MRS, WZ

Funding acquisition: CAM

Supervision: CAM

Writing – original draft: MRS, CAM

Writing – review & editing: MRS, CAM, GCT, BZ, MLS, KDK, WZ

## Data and materials availability

The RNA sequencing data used in this analysis is available through the CommonMind Consortium through an approval process.

## Supporting Information

**S1 Fig.**
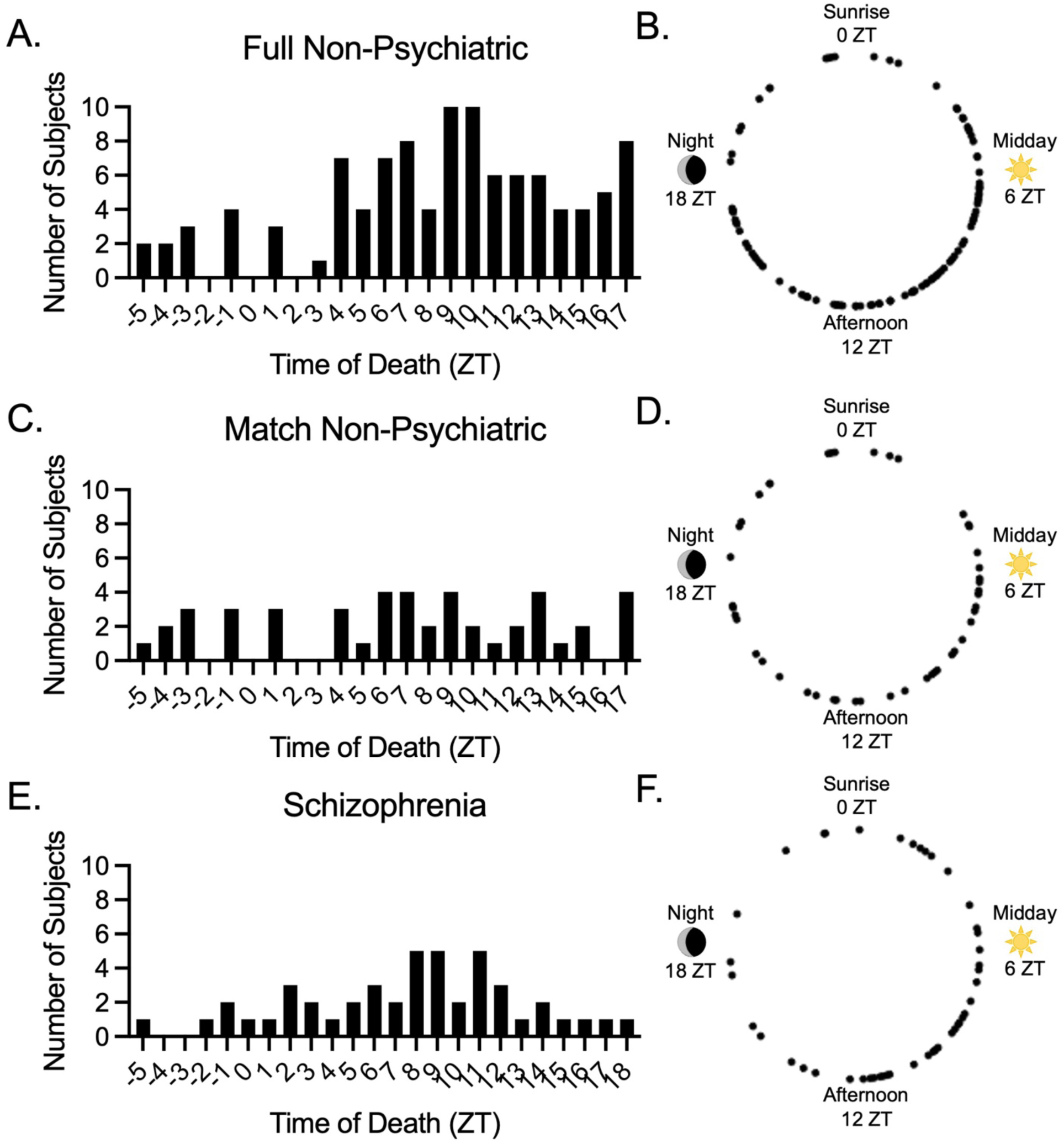
Time of death distributions. Time of Death (TOD) values for subjects in the (A-B) full no psychiatric diagnosis (*n* = 104; fNP), (C-D) match NP (*n* = 46; mNP), and (E-F) schizophrenia (*n* = 46; SZ) cohorts plotted as frequency distributions (A, C, E) and around a 24 h circle plot (B, D, F).

**S2 Fig.**
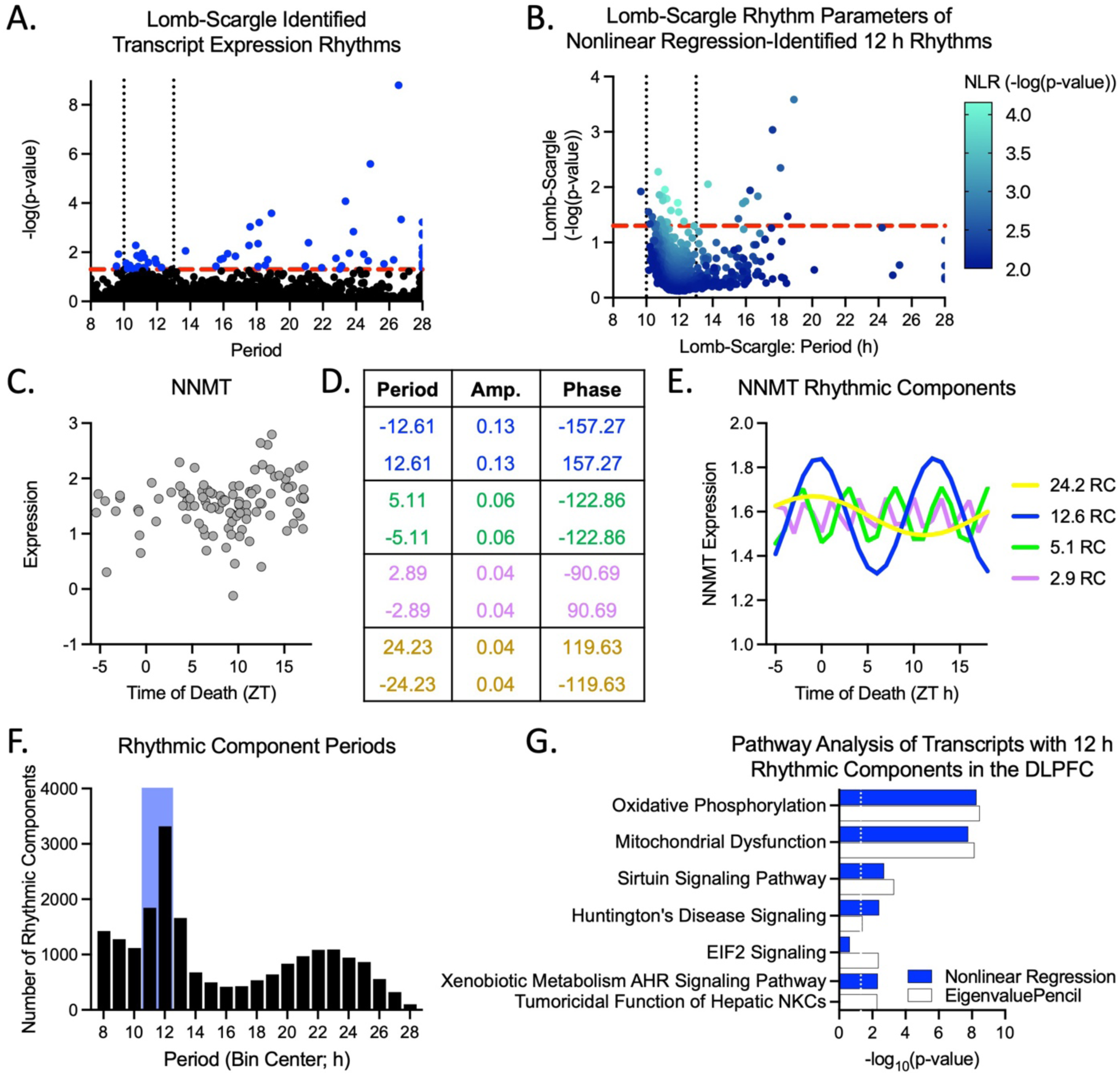
Additional rhythmicity analyses demonstrating 12 h rhythms in human DLPFC. (A-B) Lomb-Scargle analysis. Points above the red dotted lines have p-values > 0.05 in the Lomb-Scargle analysis, while black dotted lines indicate a region of period enrichment between 10-12 h. (A) Periods and p-values of all transcripts, p > 0.05 is indicated in blue. (B) The periods and p-values determined by the Lomb-Scargle analysis of transcripts identified as having a significant (p < 0.01) 12 h rhythm by the nonlinear regression (NLR) analysis. (C-E) An abbreviated example of the eigenvalue/pencil method fo a single gene is shown, with the (C) initial expression values, the (D) rhythmic components (RCs) reported by the analysis, and (E) a graphical representation of the RCs superimposed on each other. (F-G) The eigenvalue/pencil analysis in human DLPFC. (F) A histogram of RC periods, with the range of what we considered to be 12 h RCs highlighted in blue. (G) The top 5 pathways identified by an IPA of transcripts with 12 h rhythms (blue) and 12 h RCs (white).

**S3 Fig.**
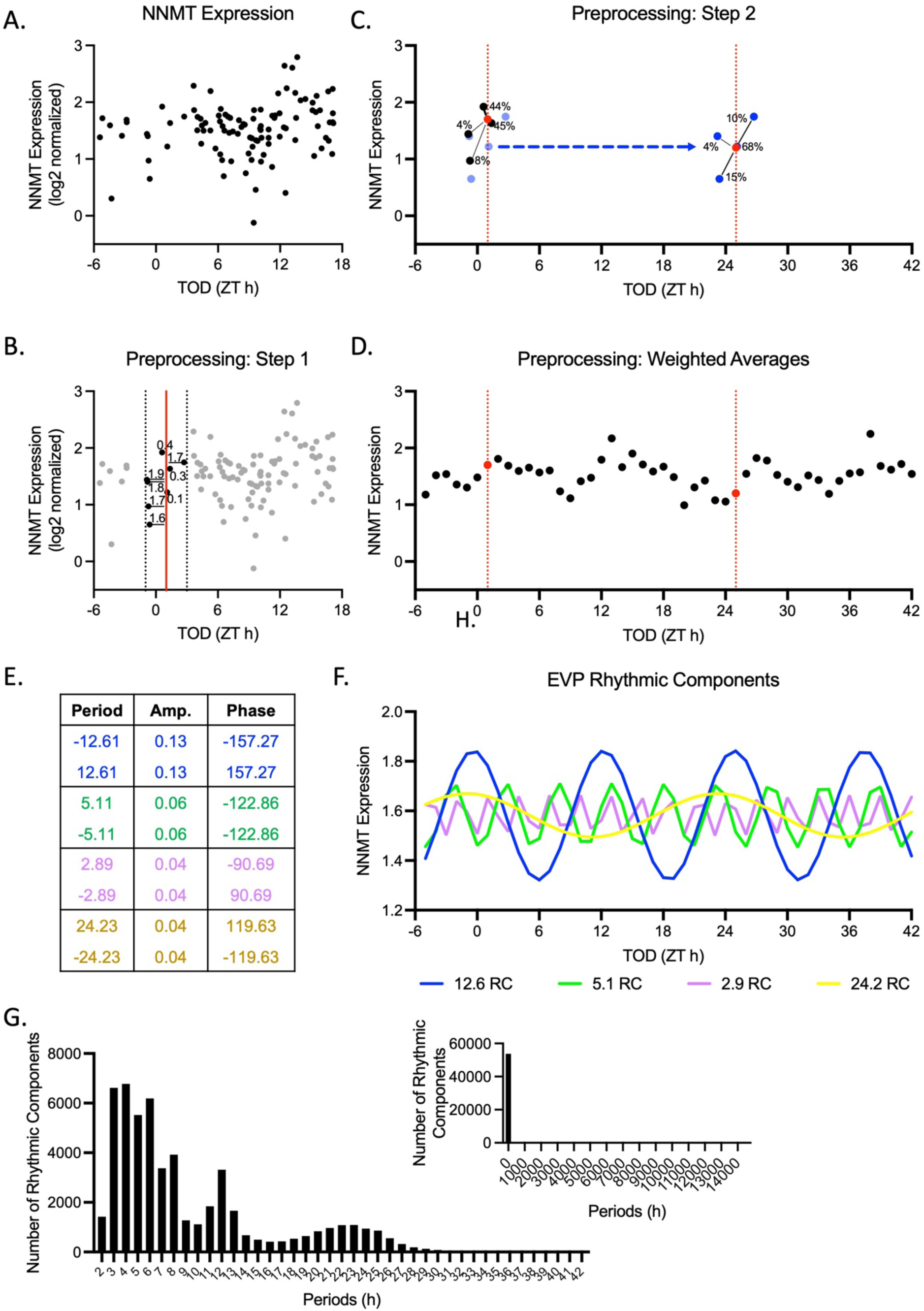
Applying the eigenvalue/pencil analysis to human postmortem brain tissue. Example of applying the eigenvalue/pencil method to RNA-seq data from human postmortem brain tissue. (A) Example of gene expression across TOD. (B-D) Preprocessing steps to convert to even-interval data. (E-F) Example of eigenvalue/pencil results for an individual gene. (G) Histogram of rhythmic component periods from all 13,915 genes analyzed.

**S4 Fig.**
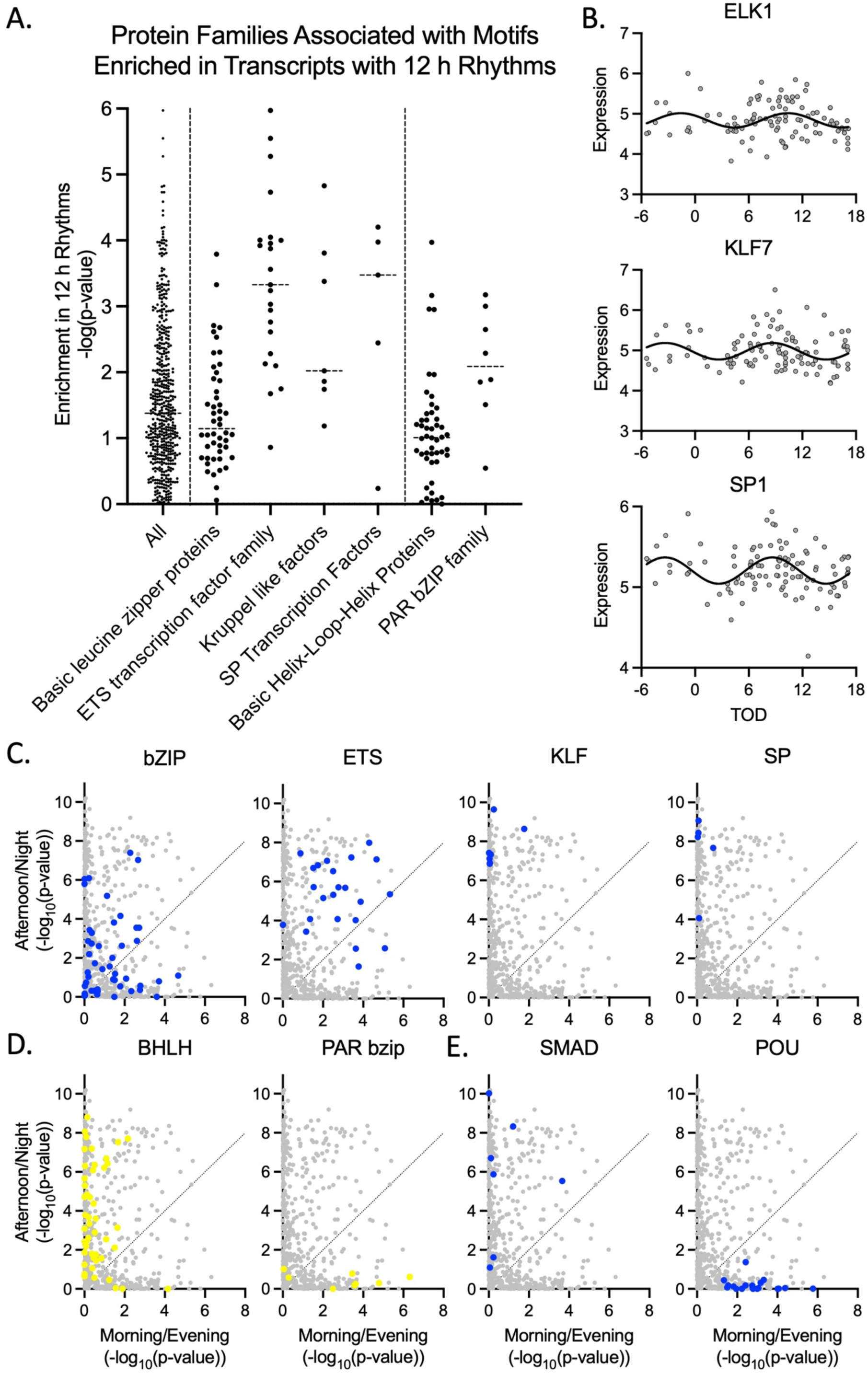
Protein families associated with motifs enriched in transcripts with 12 h rhythms. (A) Motif enrichment analysis of transcripts with 12 h rhythms in the human DLPFC. Dashed lines separate three groups: (1) All motifs tested; (2) Protein families previously implicated in regulating 12 h rhythms; (3) Protein families previously implicated in regulating 24 h rhythms. (B) Examples of transcription factors from ETS domain (Elk1), Kruppel-Like Factor (KLF7), and SP Transcription factor (SP1) families that have 12 h rhythms in human DLPFC. (C-E) Comparison of motif enrichment analysis of transcripts with 12 h rhythms that peak in the morning/evening and those that peak in the afternoon/night. (C) Protein families previously implicated in regulating 12 h rhythms. Individual motifs are marked in blue for each family. (D) Protein families previously implicated in regulating 24 h rhythms. Individual motifs are marked in yellow for each family. (E) Other top protein families associated with motif enrichment.

**S5 Fig.**
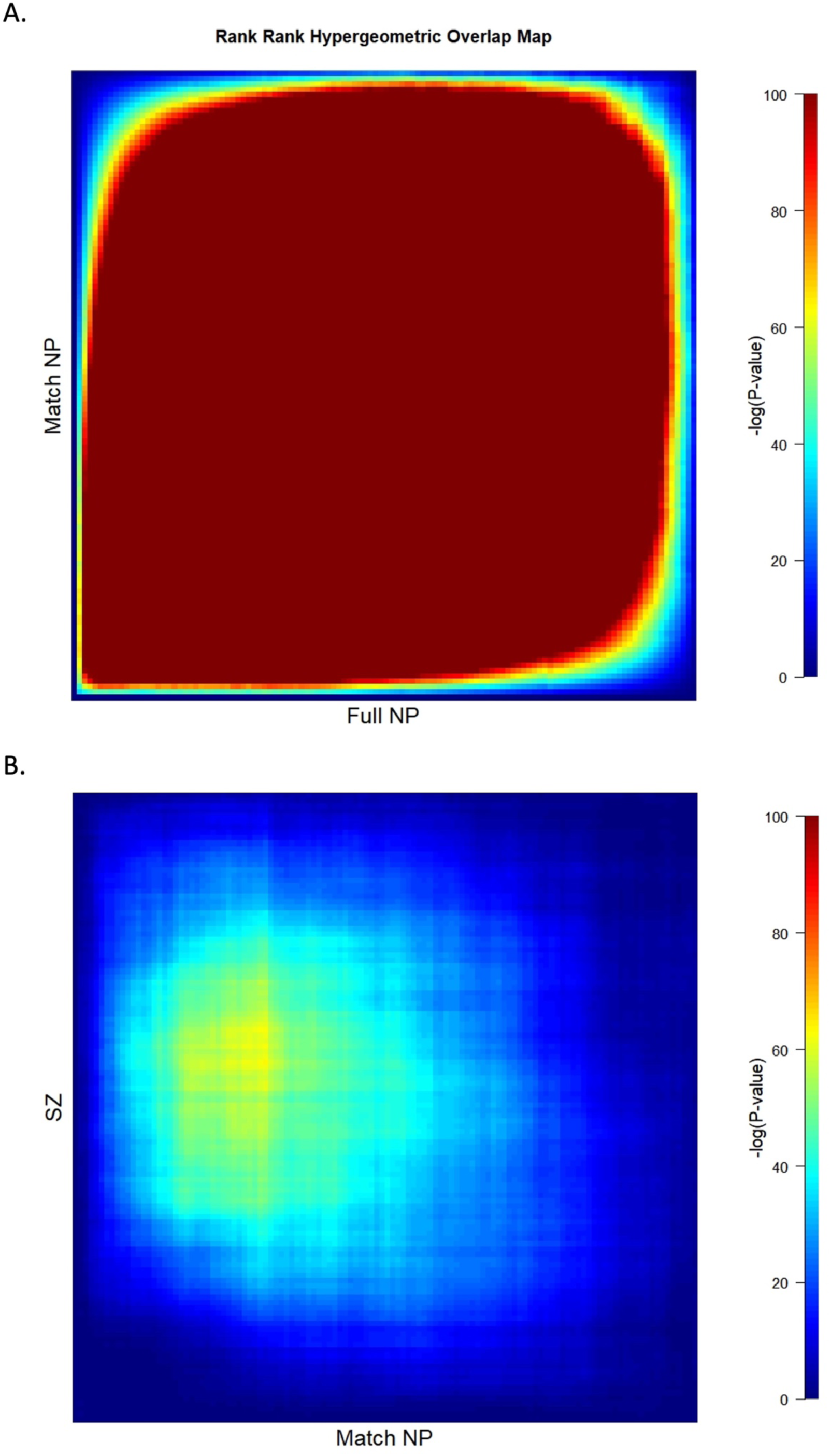
Rank-rank hypergeometric overlap comparison of 12 h rhythmicity between cohorts. RRHO plots comparing the (A) full and match NP cohorts and the (B) mNP and SZ cohorts.

**S6 Fig.**
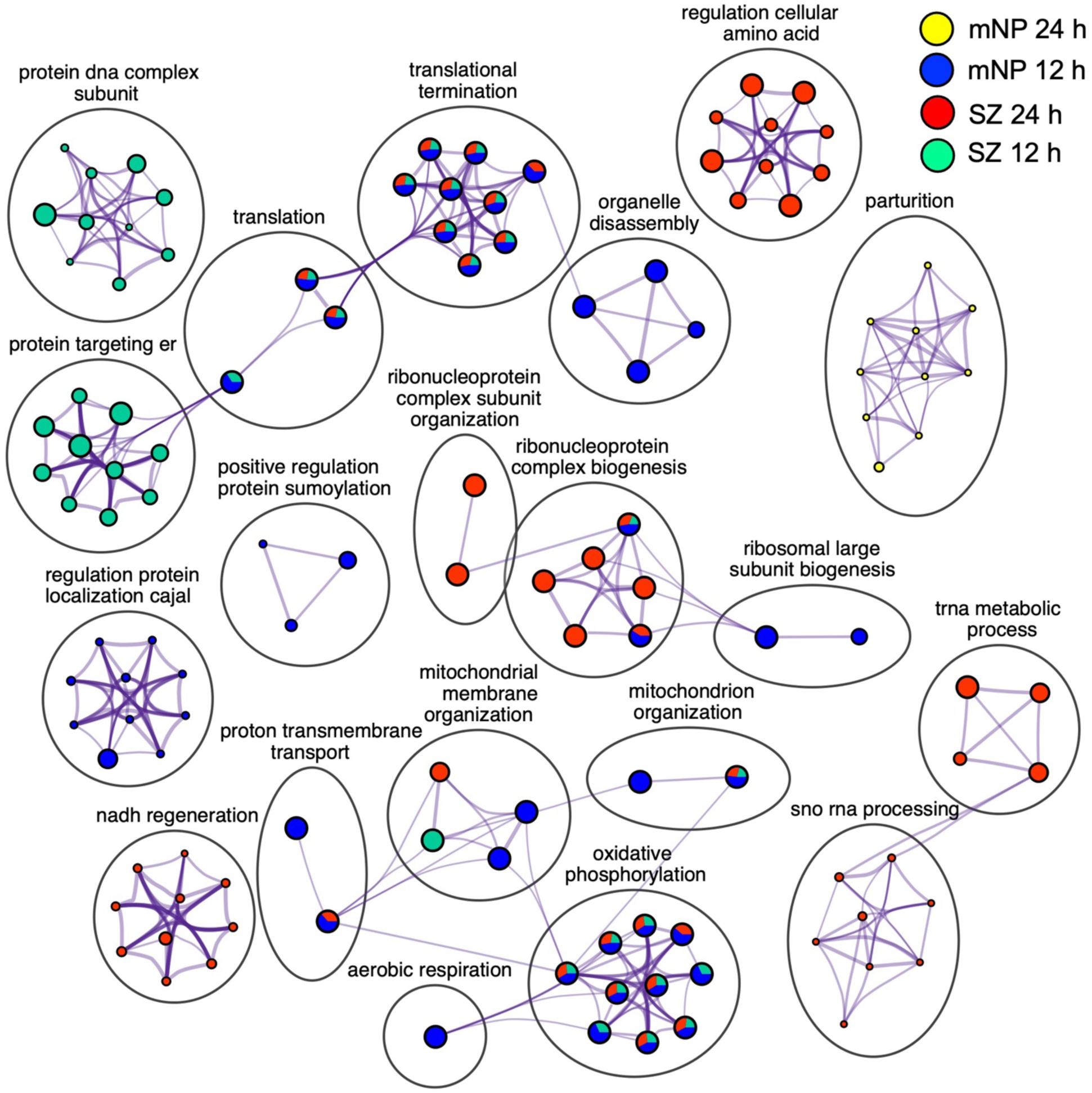
Biological processes enriched for transcripts with 12 and 24 h rhythms in expression. Cytoscape depiction of a Metascape analysis of transcripts with 12 and 24 h rhythms in both the mNP and SZ cohorts.

**S7 Fig.**
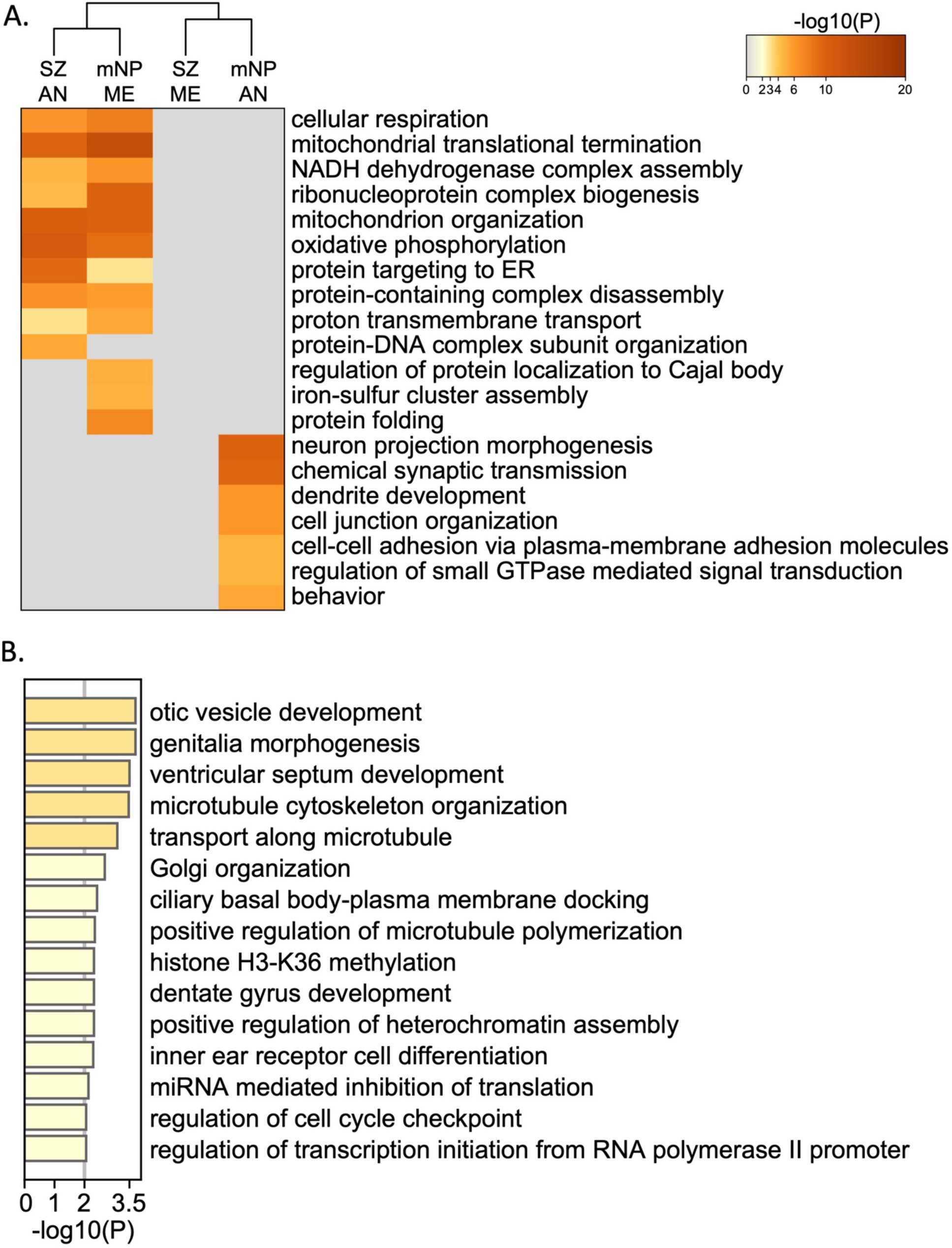
Biological processes enriched in transcripts with 12 h rhythms that peak either in the morning/evening or in the afternoon/night. (A) Heatmap of Metascape analysis of transcripts with 12 h rhythms that peak in expression either in the morning/evening (ME) or afternoon/night (AN). (B) Biological processes enriched in the SZ ME group analyzed separately due to much lower strength of enrichment than the other 3 groups.

**S8 Fig.**
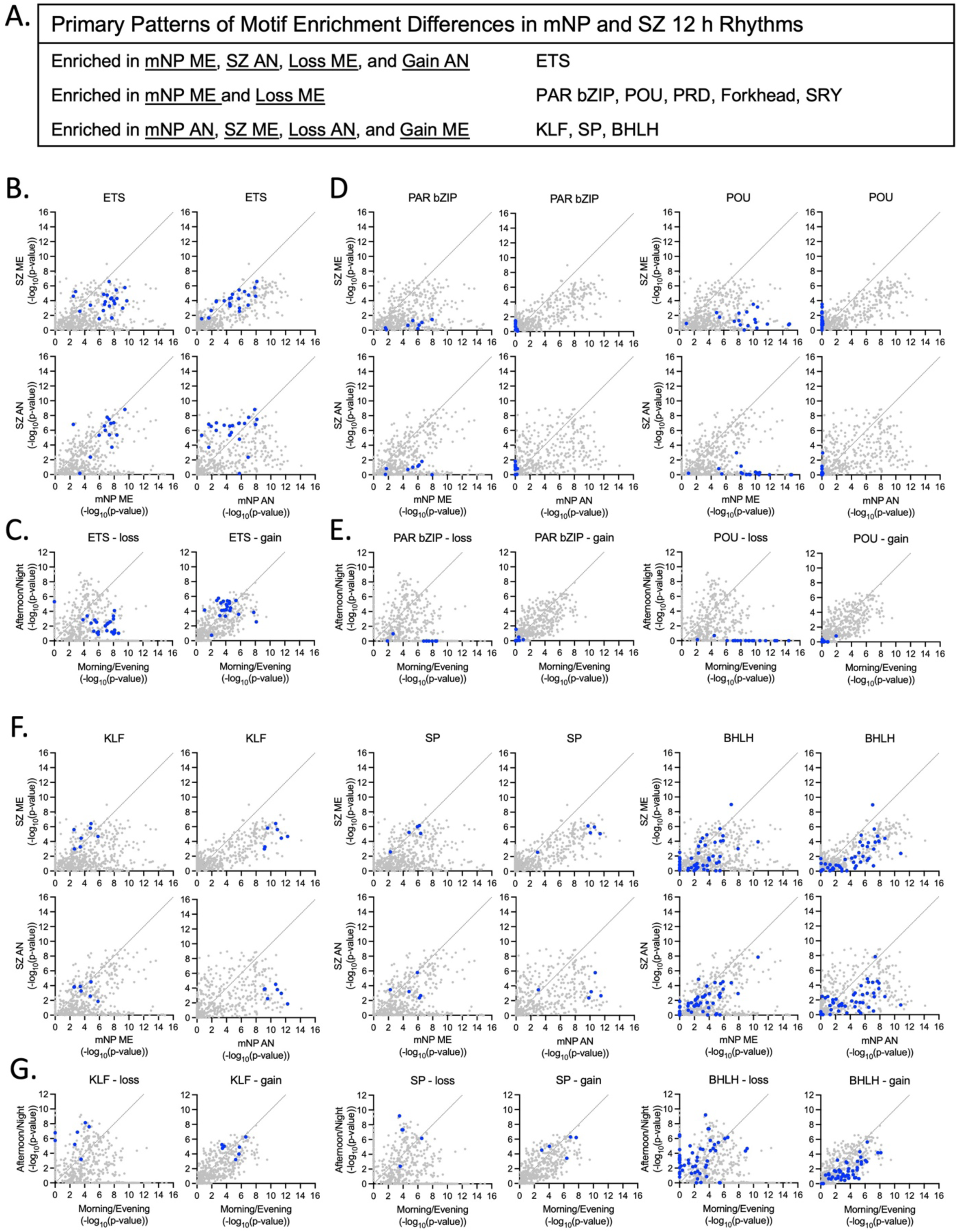
Time dependent differences in motif enrichment between mNP and SZ cohorts. (A) Summary of patterns in protein families associated that emerge after motif enrichment analysis. Families include: ETS domain family (ETS), PAR basic leucine zipper family (PAR bZIP), POU class homeoboxes, PRD class homeoboxes, Forkhead boxes, Sex determining region Y boxes (SRY), Kruppel-Like Factors (KLF), SP domain family (SP), Basic helix-link-helix domain family (BHLH). (B,D,F) Comparison of motif enrichment scores (-log10(pvalue) between mNP and SZ for transcripts that peak either in the morning/evening (ME) or afternoon/night (AN). (C,E,G) Comparison of motif enrichment scores between ME and AN groups for transcripts that either significantly lose 12 h rhythms in SZ, or significantly gain 12 h rhythms in SZ. (B-C) ETS domain family, which shows altered timing in SZ (mNP ME to SZ AN). (D-E) Examples of protein families have enrichment in mNP ME, but no enrichment in any SZ groups. (F-G) Examples of protein families with altered timing in SZ (mNP AN to SZ ME).

**S1 Table.** Description of CommonMind Consortium cohorts.

|  | Full NP (n = 104) | Match NP (n =46) | SZ (n = 46) |
| --- | --- | --- | --- |
| **Sex (Male/Female)** | 81/23 | 33/13 | 32/14 |
| **Race (White/Black)** | 86/17 | 35/11 | 34/12 |
| **Age (y, mean ± SD)** | 48.4 ± 12.3 | 49.1 ± 13.2 | 50.1 ± 11.5 |
| **PMI (h, mean ± SD)** | 17.9 ± 6.0 | 17.3 ± 6.2 | 17.0 ± 8.3 |
| **Brain pH (mean ± SD)** | 6.7 ± 0.2 | 6.6 ± 0.2 | 6.5 ± 0.3 |
| **TOD (mean ± SD)** | 8.6 ± 5.7 | 7.0 ± 6.4 | 7.8 ± 5.2 |
| **Site (Pitt/MSSM)** | 61/43 | 22/24 | 22/24 |
| **S1 Table. Description of CommonMind Consortium cohorts**<br>Abbreviations: y = years, h = hours; PMI = postmortem interval; SD = standard deviation;<br>TOD = time of death; |  |  |  |

**S2 Table.**
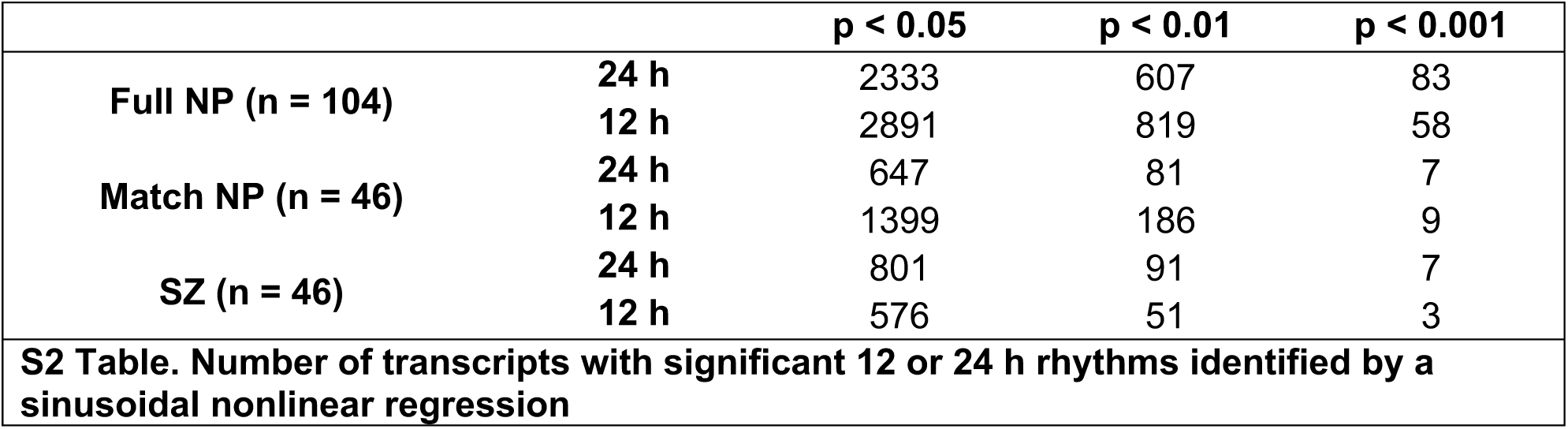
Number of transcripts with significant 12 or 24 h rhythms identified by a sinusoidal nonlinear regression.

**S1 File. Sinusoidal Nonlinear Regression Analyses**. (Tabs 1-3) Results of the sinusoidal nonlinear regression analyses identifying 12 and 24 h rhythms. (Tab 4) 12 h rhythm parameter comparisons between mNP and SZ cohorts, including gain/loss analysis.

**S2 File. Lomb Scargle Analysis**. (Tab 1) Results of the Lomb Scargle Analysis for all genes. (Tab 2) Abbreviated results of the Lomb Scargle and Nonlinear Regression Analyses for genes with significant 12 h rhythms in the Nonlinear Regression Analysis (p < 0.01).

**S3 File. Eigenvalue/Pencil Analyses**. (Tab 1) Raw eigenvalue/pencil output for all three cohorts. (Tab 2) 12 h RCs (11 ≤ Period < 13). (Tab 3) 24 h RCs (20 ≤ Period < 26). (Tab 4) Genes with both 12 and 24 h RCs.

**S4 File. Ingenuity Pathway Analyses**. Ingenuity Pathway Analyses results for all groups.

**S5 File. Metascape Analyses**. (Tab 1 – 2) Results of Metascape analysis of 12 and 24 h rhythms from mNP and SZ cohorts. (Tab 1) Top 100 enriched biological pathways. (Tab 2) Full GO list and membership results used to create cytoscape images. (Tab 3 – 4) Results of Metascape analysis of 12 h rhythms that either peak during the morning/evening or afternoon/night in the mNP and SZ cohorts. (Tab 1) Top 100 enriched biological pathways. (Tab 2) Full GO list and membership results used to create cytoscape images.

**S6 File. LISA Analyses**. (Tab 1) P-values for all motifs analyzed in LISA Combined Cistrome Motif Enrichment Analysis. (Tab 2) P-values only for motifs of genes expressed in our dataset. HGNC protein family category is listed.

